# Mechanistic Implications of Enhanced Editing by a HyperTRIBE RNA-binding protein

**DOI:** 10.1101/156828

**Authors:** Weijin Xu, Reazur Rahman, Michael Rosbash

## Abstract

We previously developed TRIBE, a method for the identification of cell-specific RNA binding protein targets. TRIBE expresses an RBP of interest fused to the catalytic domain (cd) of the RNA editing enzyme ADAR and performs Adenosine-to-Inosine editing on RNA targets of the RBP. However, target identification is limited by the low editing efficiency of the ADARcd. Here we describe HyperTRIBE, which carries a previously characterized hyperactive mutation (E488Q) of the ADARcd. HyperTRIBE identifies dramatically more editing sites, many of which are also edited by TRIBE but at a much lower editing frequency. HyperTRIBE therefore more faithfully recapitulates the known binding specificity of its RBP than TRIBE. In addition, separating RNA binding from the enhanced editing activity of the HyperTRIBE ADAR catalytic domain sheds light on the mechanism of ADARcd editing as well as the enhanced activity of the HyperADARcd.

## Introduction

Pre-mRNAs and mRNAs are subject to many post-transcriptional regulatory events, which are mediated by RNA-binding proteins (RBPs). RBPs have been found to be essential for pre-mRNA splicing, 3’ end formation, mRNA translocation from the nucleus to the cytoplasm, and translation among other processes (Zhao et al. 1999; Jansen 2001; Witten and Ule 2011; Szostak and Gebauer 2013; Gerstberger et al. 2014). Numerous human diseases have been linked to RBPs, e.g., Amyotrophic Lateral Sclerosis (ALS) (Nussbacher et al. 2015). Identifying the RNA targets of a RBP is therefore not only crucial for deciphering its function but also an important aspect of deciphering RBP-related human diseases.

The current standard technique for identifying RBP targets *in vivo* is CLIP (Cross-Linking and ImmunoPrecipitation) and its variants (Ule et al. 2005; Hafner et al. 2010). These methods use UV light to covalently link the RBP to its targets, immunoprecipitate the RNA-protein complex, and then purify and identify the covalently bound RNA. CLIP has definite advantages, for example the ability to identify the exact RBP binding sites on RNA, but there is also a disadvantage: its requirement for large amounts of material to make an extract and do immunoprecipitations (Darnell 2010). It is therefore often difficult to perform CLIP in a cell-specific manner.

We recently developed TRIBE (Targets of RNA-binding proteins Identified By Editing) to study RBP targets in specific cells, especially in small numbers of circadian neurons of the *Drosophila* adult brain. This method expresses in specific cells a fusion protein of an RBP and the catalytic domain of *Drosophila* ADAR (McMahon et al. 2016). ADAR is a well-conserved RNA editing enzyme. It deaminates adenosine to inosine, which is read by ribosomes and reverse transcriptase as guanosine (Basilio et al. 1962; Bass and Weintraub 1988). ADAR consists of two modular parts, double-strand RNA-binding motifs (dsRBMs) and a catalytic domain (Kim et al. 1994; O'Connell et al. 1995; Macbeth et al. 2005). TRIBE replaces the dsRBMs of the *Drosophila* ADAR enzyme with the RBP of interest (McMahon et al. 2016). Fusion protein binding and mRNA specificity are therefore driven by the RBP due to the absence of the dsRBMs. TRIBE-dependent editing sites are identified by deep sequencing of RNA extracted from cells of interest. The editing percentage of any mRNA nucleotide is the number of adenosines edited to inosines divided by total number of reads at that site. In the initial TRIBE paper (McMahon et al. 2016), we used a conservative criterion to minimize false positives, namely, a site is only counted if it has an editing percentage greater than 10%, and a read coverage greater than 20, i.e., at least 2 editing events/20 reads in both biological replicates.

Although we were successful in identifying bona fide targets of three RBPs (the *Drosophila* HnRNP protein Hrp48, the nuclear protein NonA and FMRP, the *Drosophila* version of the mammalian Fragile X protein), the number of targets identified by TRIBE in tissue culture was substantially reduced compared to CLIP data with the same RBP (McMahon et al. 2016). Even assuming some CLIP false positives (Darnell 2010; Lambert et al. 2014), TRIBE may still be experiencing a high false-negative problem. This could be due in part to innate editing specificity of the ADARcd. It preferentially edits adenosines bordered by 5’ uridines and 3’ guanosines, i.e., a UAG sequence (Lehmann and Bass 2000; Eggington et al. 2011), which is also observed in TRIBE (McMahon et al. 2016). The ADARcd also prefers to edit adenosines surrounded by a double-stranded region (Bass and Weintraub 1988; Kim et al. 1994; O'Connell et al. 1995; Eggington et al. 2011; Matthews et al. 2016). Consequently, a substantial fraction of RBP-target mRNAs may go unidentified.

To enhance the efficiency and/or reduce the specificity of the ADARcd, we turned our attention to a mutational screen directed at the catalytic domain of human ADAR2 (Kuttan and Bass 2012). It identified a “hyperactive” E488Q mutation, which was reported to have these precise characteristics. Ideally, incorporating this hyperactive E488Q mutation into TRIBE (HyperTRIBE) could reduce its false negative rate.

Hrp48 HyperTRIBE, which carries this E488Q mutation within the *Drosophila* ADARcd (dADARcd), indeed identifies dramatically more editing sites than Hrp48 TRIBE. Many of these sites correspond to below-threshold TRIBE editing targets. This indicates that they are bona fide targets, which are edited more efficiently by HyperTRIBE. They also overlap much more successfully with CLIP data, indicating that HyperTRIBE has a much reduced false-negative problem and suggesting that it more faithfully recapitulates the known binding specificity of its RBP than TRIBE. The HyperTRIBE data also have mechanistic implications about the TRIBE fusion protein strategy as well as about the function of the ADARcd as well as its E488Q variant.

## Results

The original Hrp48 TRIBE construct was made by fusing coding DNA for the Hrp48 protein followed by a short linker and then dADARcd (henceforth called TRIBE). To make the companion HyperTRIBE construct, we first identified the dADARcd glutamate corresponding to human ADAR2 E488. That residue as well as surrounding sequence is highly conserved between hADAR2 and the dADARcd (data not shown), suggesting that it should function similarly in the *Drosophila* enzyme. We then introduced the E488Q mutation into the original Hrp48 TRIBE construct via QuikChange^®^ site-directed mutagenesis. We had no difficulty making a stable S2 cell line expressing Hrp48 TRIBE (McMahon et al. 2016) but failed with Hrp48 HyperTRIBE (henceforth called HyperTRIBE), perhaps because of the greatly enhanced editing frequency (see below). Most experiments were therefore performed by transiently expressing fusion proteins in *Drosophila* S2 cells together with GFP and sorting GFP-positive cells by FACS. Editing sites were always defined as the sites in common between two experiments, i.e., >10% editing and > 20 reads/nucleotide in both biological replicates (McMahon et al. 2016). A third replicate of TRIBE minimally reduced the number of common sites, i.e., 80% were present in the third replicate (data not shown). Transient and stable expression of TRIBE in S2 cells detected comparable target gene numbers (200-300), ~40% of which were identical (Fig. S1). This rather low rate of overlap is likely due to the substantial difference between transient and stably expressing cells.

Expression of HyperTRIBE resulted in approximately 20X the number of editing events compared to TRIBE (Fig. 1A). Expression of the ADARcd with E488Q mutation alone does not increase the number of editing events above the endogenous level of S2 cells (Fig. 1A). This is despite the fact that the HyperADARcd is stable and expressed at comparable levels to those of the other TRIBE constructs (data not shown), so most if not all editing by HyperTRIBE -- like editing by regular TRIBE -- requires the RNA binding ability of the fused RBP (McMahon et al. 2016).

**Figure 1.**
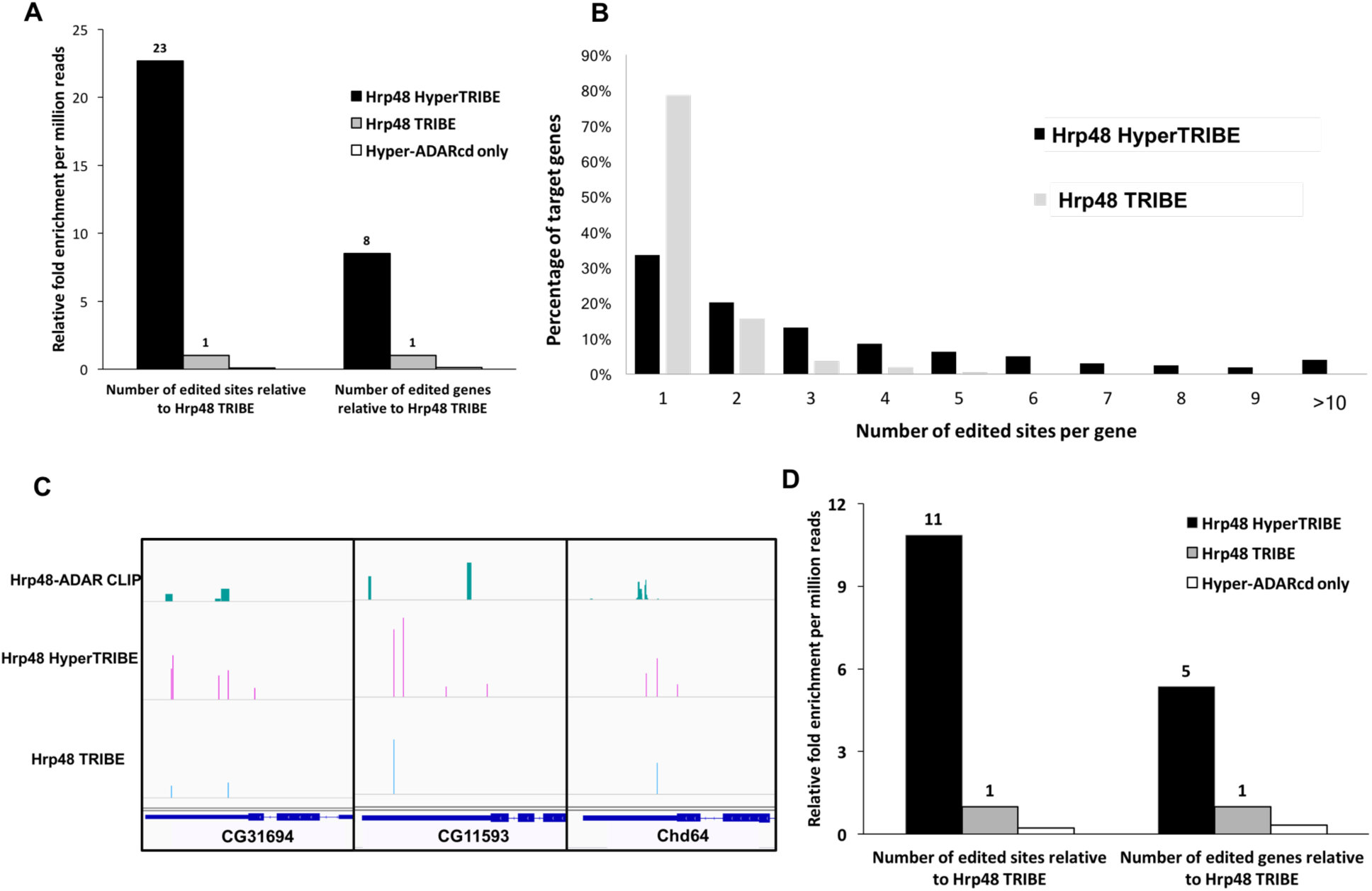
HyperTRIBE dramatically increases both the number of target genes and the number of edited sites compared to TRIBE. (A) Although HyperTRIBE and TRIBE increases in both the number of editing events and genes edited in S2 cells, the increases are much more dramatic in HyperTRIBE-expressing cells. Numerically, HyperTRIBE identifies 10689 common edited sites in two replicates, whereas the corresponding number for Hrp48 TRIBE is 291. Both TRIBE constructs have a reproducibility of around 60% and only replicable editing events are reported. There is no comparable increase in editing sites or genes with expression of the Hyper-ADARcd is alone (11 sites identified). The number of genes and by relative fold change compared to TRIBE (see also Methods). (B) A much larger fraction of target genes are edited at multiple sites by HyperTRIBE than by TRIBE. The histogram indicates the percentage of target genes containing 1 to more than 10 editing sites. Genes with multiple sites may be transcripts bound more stably by Hrp48. (C) Three examples of commonly identified genes by HyperTRIBE, TRIBE and CLIP are shown in the IGV genome browser. Hrp48-ADARcd CLIP data is from a previous publication (McMahon et al. 2016). Typically, the multiple editing sites in HyperTRIBE cluster nearby the original sites identified by TRIBE. The height of the bars indicates editing frequency for TRIBE data and CLIP signal strength for CLIP data. (D) Expression of HyperTRIBE in all neurons (with the elav-gsg-Gal4 driver) in fly brains also increases the number of editing sites and genes compared to TRIBE. Editing events are identified and normalized as in (A).

The ratio of HyperTRIBE-edited genes compared to TRIBE-edited genes is only 8 (Fig. 1A), 3-fold lower than the editing site ratio, indicating a substantial increase in the number of edited sites per gene in HyperTRIBE. Indeed, HyperTRIBE generates many more multiple-edited genes with a median of 3 edited sites per gene comparing to a median of 1 for TRIBE (Fig. 1B). Some HyperTRIBE editing sites are near the original TRIBE sites (Fig. 1C), suggesting that the higher editing rate of HyperTRIBE is due in part to its ability to edit multiple adenosines near the original ADARcd interacting region. Moreover, the data show that HyperTRIBE unique editing sites that are on the same molecule as common sites – within a single RNA-seq read – have a higher editing percentage than all unique editing sites (Fig. S2). This indicates that the HyperADARcd may edit additional, nearby adenosines without fully releasing the mRNA.

To extend the enhanced editing of HyperTRIBE to *Drosophila,* we performed cell-sorting experiments on flies expressing HyperTRIBE in all adult fly brain neurons using the elav-gsg-Gal4 driver (Abruzzi et al. 2015; McMahon et al. 2016). Similar to the tissue culture result, HyperTRIBE exhibits a 11-fold increase in the number of edited sites and a 5-fold increase in the number of edited genes compared to TRIBE (Fig. 1D).

We next compared the HyperTRIBE editing data with our previous CLIP data as well as with regular TRIBE editing data in *Drosophila* S2 cells. HyperTRIBE not only identifies 282 (97%) of the edited sites and 220 (98%) of the edited genes identified by TRIBE (Fig. 2A, 2B), but these data also correlate well with the CLIP results: 73% of the HyperTRIBE sites are identified by CLIP. (78% for the TRIBE sites; Fig. 2B). However, there is a striking difference of 66% vs 4% in the ability of HyperTRIBE and TRIBE to recognize CLIP-identified genes, respectively (Fig. 2B and S3). This distinction is due to the many fewer sites and genes identified by TRIBE, indicating that HyperTRIBE significantly lowers the TRIBE false negative rate and thereby provides a much more complete binding signature of the RBP. We obtained similar results with FMRP HyperTRIBE (Fig. S4), another RBP assayed in the original TRIBE paper, indicating that the higher efficiency of HyperTRIBE is not limited to Hrp48.

**Figure 2.**
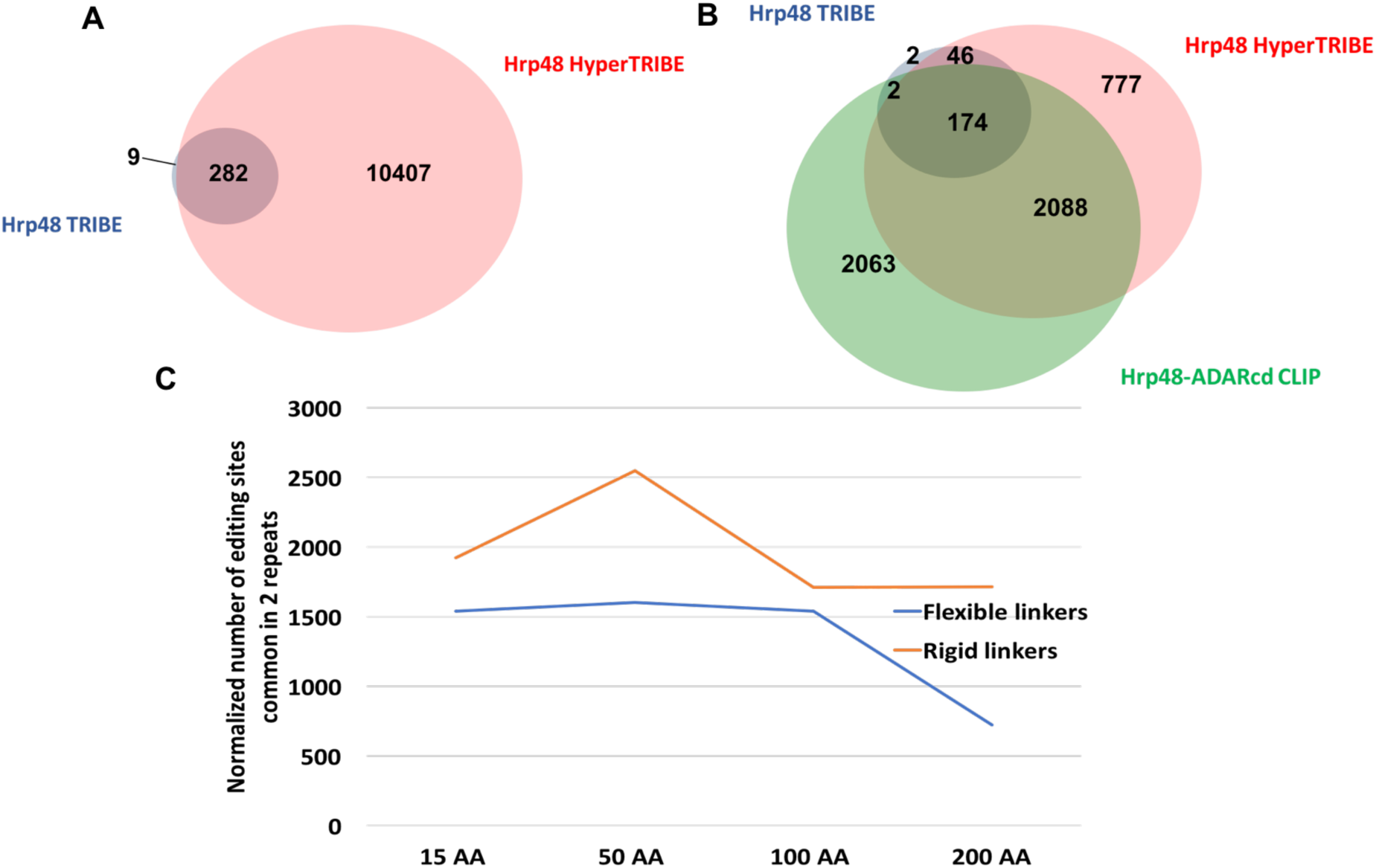
HyperTRIBE data faithfully reflect Hrp48 binding specificity with higher sensitivity than TRIBE. (A) Venn diagram of the editing sites shows that almost all of the Hrp48 TRIBE sites (blue) are also detected by HyperTRIBE (pink). (B) Both TRIBE- and HyperTRIBE-identified genes overlap well with CLIP-identified genes. The Venn diagram shows the overlap of all genes identified by TRIBE (224 in total, blue), HyperTRIBE (3085 in total, pink) and Hrp48-ADARcd CLIP (4327 in total, green) (McMahon et al. 2016). The overlap between TRIBE and CLIP is not significantly different from the overlap between HyperTRIBE and CLIP (Z-test performed, p=0.09). (C) Editing efficiency of HyperTRIBE decreases with linker length. Flexible linkers are repeats of (GGGGS)_n_ of 15 AA, 50 AA, 100 AA and 200 AA length (blue), and rigid linkers are repeats of (EAAAK)_n_ of 15 AA, 50 AA, 81 AA and 162 AA length (yellow). Minor modifications were made to the peptide sequence for cloning convenience, by substituting amino acids of similar properties. Editing sites are identified and normalized as indicated in Fig. 1A.

Because only a minimal seven amino acid linker was used between Hrp48 and the ADARcd in the original TRIBE paper (McMahon et al. 2016) and in the HyperTRIBE assays shown to this point, we investigated whether expanding the linker and altering its character would impact HyperTRIBE editing. We imagined that a longer and more flexible linker might increase editing, perhaps even dramatically, or it might decrease editing if proximity to the target RNA was important. Editing efficiency is indeed decreased but only about two-fold, even with a 200aa flexible linker (Fig. 2C). This indicates that the precise relationship between the RBP and the ADARcd is not critical, also suggesting that other means of delivering the ADARcd to RNA should be possible, for example via protein dimerization schemes (Stankunas et al. 2003).

To further address possible reasons for the additional HyperTRIBE editing, we compared the editing frequencies of the sites identified by both HyperTRIBE and TRIBE. We divided the HyperTRIBE editing sites into two categories, unique sites and common sites; the latter were also identified by TRIBE. Although HyperTRIBE edits its unique sites at similar frequencies to the much smaller number of TRIBE-edited sites, the common sites experience much higher editing frequencies with HyperTRIBE than with TRIBE (Fig. 3A). This is true not only on average but also when the sites are examined individually (Fig. 3B). This indicates that common sites are also preferred HyperTRIBE substrates.

**Figure 3.**
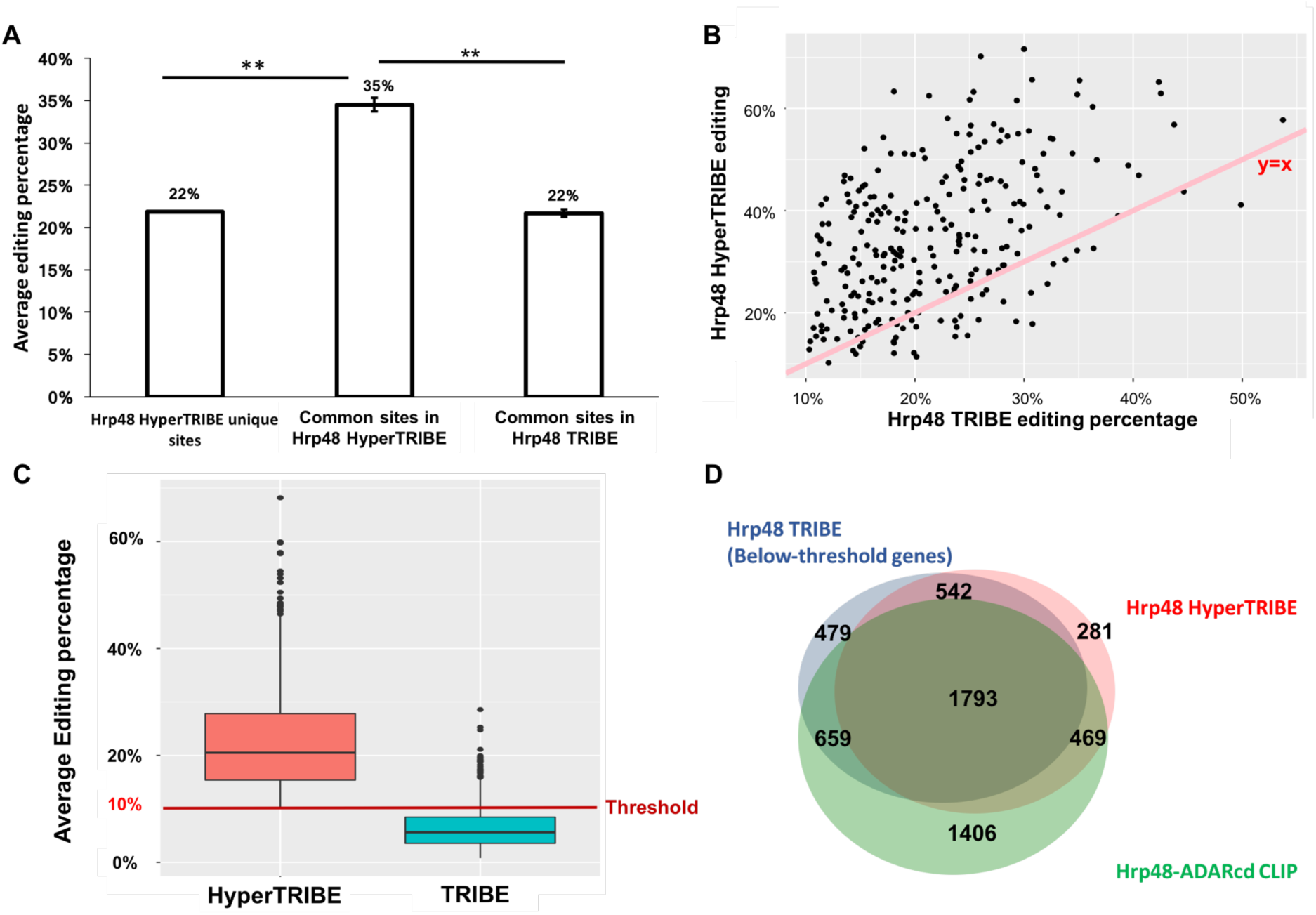
HyperTRIBE increases the editing frequency, which allows the detection of many below-threshold TRIBE sites. (A) Bar graph shows the editing percentage of the sites uniquely identified in HyperTRIBE as well as the editing percentage of the common editing sites in HyperTRIBE and in TRIBE, respectively. Editing percentage is shown as the weighted average. The editing percentage of the sites commonly identified by TRIBE and HyperTRIBE is increased in HyperTRIBE (Student t-test performed, **p<0.0001). (B) An increase in editing percentage by HyperTRIBE is observed for most sites. A scatter plot shows editing percentage of HyperTRIBE editing sites (Y-axis) and TRIBE editing represents y=x is shown for reference (pink). (C) About 30% of the 4017 below-threshold sites in TRIBE show an elevated editing percentage in HyperTRIBE, allowing them to be identified in the analysis pipeline (Z-test performed to test enrichment, P«0). HyperTRIBE editing sites that are present in TRIBE but below the 10% threshold are shown. Some below-threshold TRIBE editing sites have > 10% average editing percentage because one replicate has >10% editing but the other replicate has <10%. A red line indicating 10% threshold is shown for reference. (D) 67% and 71% of TRIBE below-threshold target genes overlap with HyperTRIBE and CLIP respectively. Venn diagram shows the overlap of genes edited below-threshold by TRIBE (3473 in total, blue), genes identified by HyperTRIBE (3085 in total, pink) and Hrp48-ADARcd CLIP genes (4327 in total, green) (McMahon et al. 2016). Although small, the difference in overlap between below-threshold TRIBE genes and CLIP (71%) versus overlap between HyperTRIBE genes and CLIP (73%) is significant (Z-test performed, p=0.015).

Is it possible that the unique sites are also quantitatively rather than qualitatively different between HyperTRIBE and TRIBE? This suggests that many of them might be edited by TRIBE but below the required threshold. Indeed, there are 4017 adenosines that meet this criterion, i.e., at least one editing event for each specific adenosine in both replicates but with less than 10% editing percentage. The correspondence between replicates for these adenosines and HyperTRIBE editing sites is highly significant (Z-test performed, p-value≈0), and more than 30% of these 4017 adenosines correspond to HyperTRIBE editing sites (Fig. 3C). This number is much greater than the number of C-to-T editing/mutations, which serve as control events (data not shown). Notably, these 4017 below-threshold editing sites occur in 3473 different genes, which overlap well with genes identified by Hrp48 HyperTRIBE and Hrp48-ADARcd CLIP (Fig. 3D). We conclude that below-threshold TRIBE-edited adenosines make a significant contribution to the extra HyperTRIBE editing sites and that much of the distinction between HyperTRIBE and TRIBE is quantitative.

The ADARcd with the hyperactive mutation (E488Q) has been shown to have less neighboring sequence preference surrounding the edited adenosine (Kuttan and Bass 2012). Not surprisingly perhaps, this preference for 5’ uridines and 3’ guanosines is also reduced for the sites identified by HyperTRIBE compared to TRIBE (Fig. 4A). This result indicates that much if not most of this neighboring sequence preference derives from the ADARcd, despite an indication that the 3’ guanosine preference might derive from the ADAR dsRBDs (Stefl et al. 2010). Although binding could be sequential, i.e., first dsRBD recognition followed by a conformational change and ADARcd recognition, the ADARcd appears to contain most of the specificity (Matthews et al. 2016).

**Figure 4.**
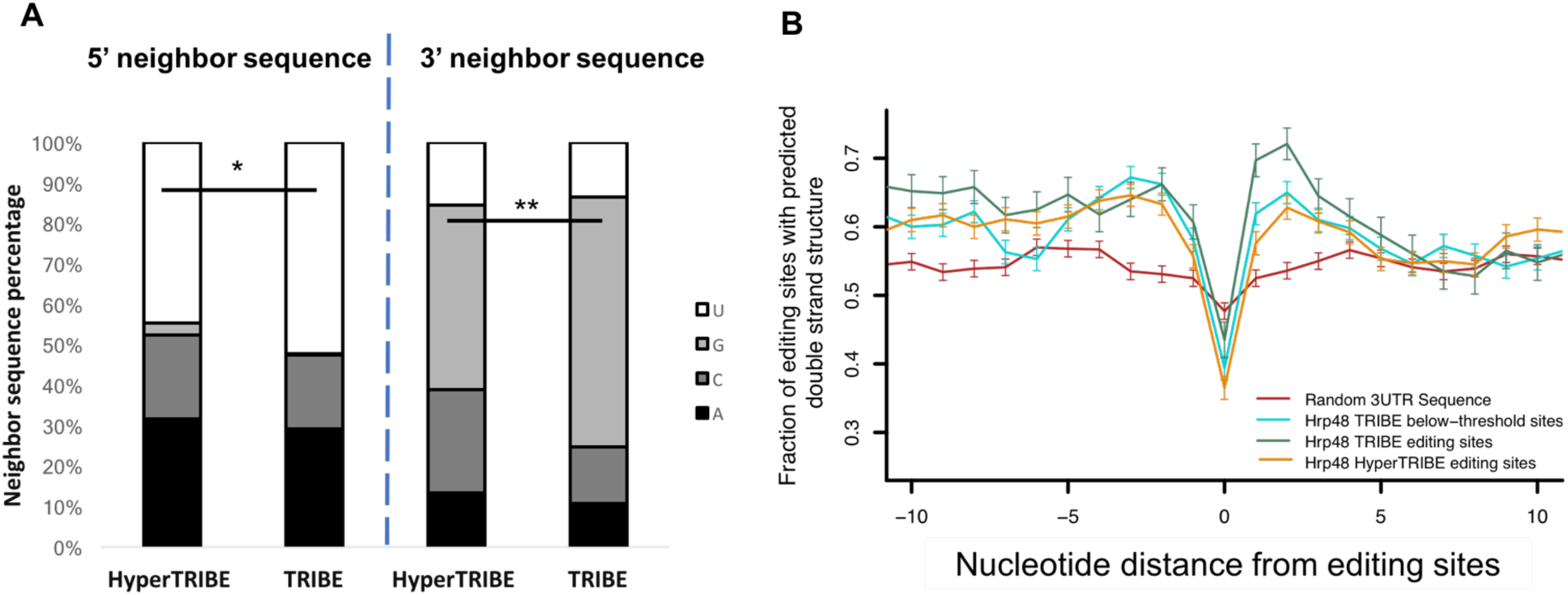
HyperTRIBE has less nearest neighbor sequence and double-stranded structure requirements than TRIBE. (A) 5’ and 3’ immediate neighbor sequence preference is reduced in HyperTRIBE. ADAR shows a neighbor preference of uridine at the 5’ and guanosine at the 3’ side. This preference is lower in HyperTRIBE than in TRIBE (Z-test performed, *p<0.05, **p<0.0001, percentage of uridine in 5’ neighbor and percentage of guanosine in 3’ neighbor tested, HyperTRIBE vs TRIBE). (B) Predicted double-strandedness is comparable surrounding TRIBE below-threshold sites and HyperTRIBE editing sites, both of which are lower than TRIBE editing sites. However, all of them have higher preference for double-stranded structure than random 3’ UTR sequences. The dip at the edited adenosines corresponds to the preference of ADARcd for bulged adenosines as substrates, which is less prominent in the random 3’ UTR sequences (centered on an adenosine). Double-strandedness was examined with a custom pipeline that uses UNAFold to fold RNA sequence *in silico,* random 3’ UTR sequence (red), TRIBE below-threshold sites (turquoise), TRIBE editing sites (green) and HyperTRIBE editing sites (orange) and plotted as fraction of sites with predicted double-strandedness (see also Methods). 1000 sites were blindly selected from each pool (N=221 for Hrp48 TRIBE) as the input for this analysis. X-axis represents the relative distance from the editing sites, and Y-axis indicates the average predicated double-strandedness for each site. Error bars show the standard error.

We also calculated the preference of TRIBE and HyperTRIBE for local double-stranded structure surrounding the edited adenosines. Based on the preference of Hrp48 for mRNA 3’UTR regions, random 3’UTR sequences of the same length and centered on an adenosine were used as a control. This strategy was similar to that used previously for TRIBE sites (McMahon et al. 2016), but many more sites were included in this calculation.

As previously observed for TRIBE (McMahon et al. 2016), there is a strong HyperTRIBE preference for double-stranded structure surrounding a relatively unstructured (bulged) adenosine, features that are absent from the control 3’UTR sequences (Fig. 4B). Notably, the 3’ side of the TRIBE editing sites was more structured than the 5’ side. This asymmetry was not present in the HyperTRIBE sites, indicating a less stringent requirement of HyperTRIBE for substrate structure on the 3’ side of the editing sites (see Discussion). Not surprisingly, the below-threshold sites appear similar to the HyperTRIBE sites (Fig. 4B) despite the higher HyperTRIBE editing frequency.

To address the edited sites in different mRNA regions, we compared the fraction of sites between coding, 5’UTR and 3’UTR sequences. This distribution is an indication of RBP specificity since Hrp48 preferentially binds to mRNA 3’UTR regions with CLIP as well as with TRIBE assays (McMahon et al. 2016). When normalized for read coverage in the different mRNA regions, HyperTRIBE still preferentially edits 3’UTR sites but with a 3.5-fold preference, somewhat less than the 5-fold preference of TRIBE (Fig. S5). Moreover, coding sequence (CDS) editing preference has increased in HyperTRIBE compared to TRIBE, from 0.4 to 0.7. This is probably due at least in part to the enhanced ability of HyperTRIBE to edit more efficiently adenosines further from the RBP binding sites.

To address this issue directly, we analyzed the distance of the edited sites from the Hrp48-ADARcd CLIP peaks. They presumably locate the positions of RBP binding. The HyperTRIBE edited sites are indeed located further away from the CLIP peaks comparing to the TRIBE sites (Fig. 5A). For example, HyperTRIBE has only 40% of its sites located within 100 nt of the CLIP peaks versus 60% for TRIBE, whereas the fraction of edited sites further than 500 nt from the CLIP peaks is 29% and 15%, respectively (Fig. 5A). Presumably, more of the distant sites are poorly edited by TRIBE and then fall below the required threshold (see below). Not surprisingly perhaps, these below-threshold TRIBE sites are also located further from the CLIP peaks than regular (above-threshold) TRIBE sites (Fig. 5A). Proximity presumably contributes to a higher editing frequency.

**Figure 5.**
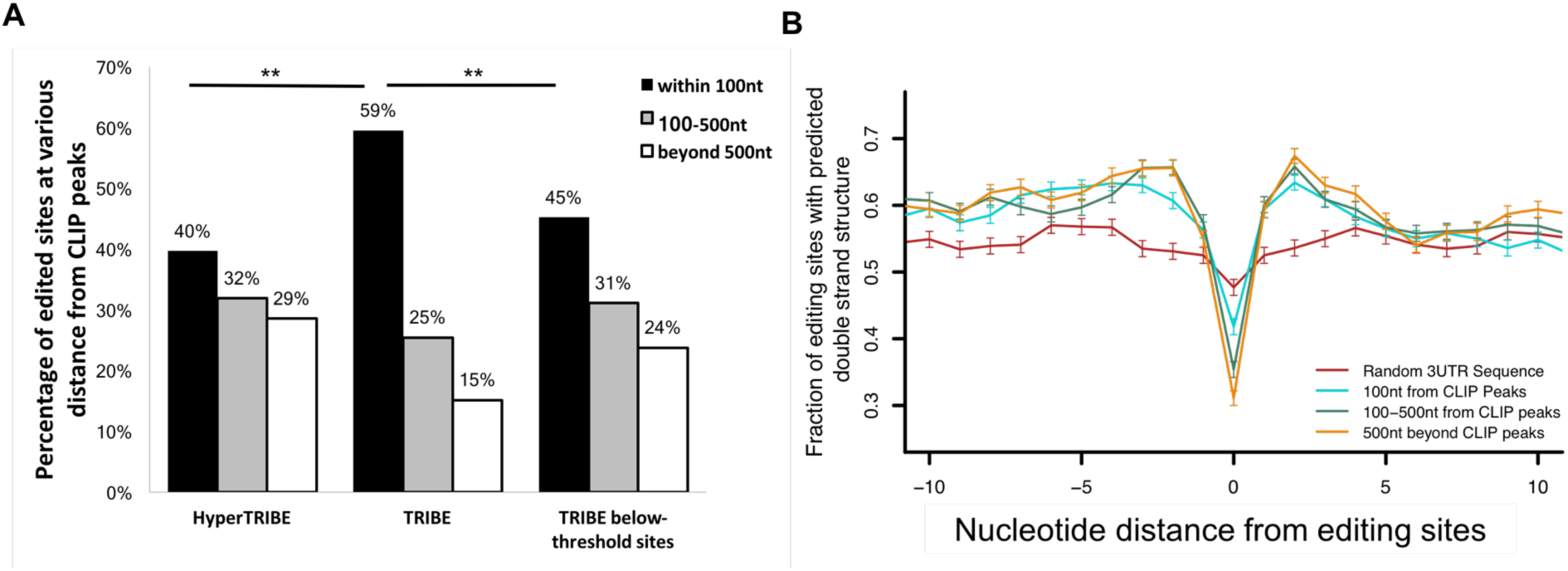
HyperTRIBE edits distant adenosines more efficiently than TRIBE. (A) Bar graph showing the distribution of editing sites relative to Hrp48-ADARcd CLIP peaks. Editing sites are categorized into three classes according to their distance from Hrp48-ADARcd CLIP peaks: within 100 nt of CLIP peaks, between 100-500 nt and greater than 500 nt (McMahon et al. 2016). A smaller fraction of editing sites in HyperTRIBE are within 100 nt of CLIP peaks (40% versus 59%). (Z-test performed, **p <0.0001, HyperTRIBE compared with TRIBE.) The below-threshold TRIBE sites show a similar pattern as HyperTRIBE sites and are more distant than above-threshold TRIBE sites. (Z-test performed, **p <0.0001, TRIBE editing sites compared to TRIBE below-threshold sites). (B) Editing sites that are closer to Hrp48 binding sites (CLIP peaks) have less local structure requirement compared to more distant sites. HyperTRIBE editing sites are categorized by their distance to Hrp48-ADARcd CLIP peaks as described in Fig. 5A. Sites within 100 nt of CLIP peaks (turquoise) show less flanking double-stranded structure and a less pronounced bulge at the edited adenosines than the 100-500 nt group (green) and beyond 500 nt group (orange). However, all three groups exhibit more surrounding double-stranded structure and a more pronounced bulged adenosine than random 3’ UTR sequence, indicating the presence of a local structural preference even for the clos editing sites. 1000 editing sites were randomly selected from each group as the input for folding analysis (see Methods).

A structural comparison between edited sites at different distances from the CLIP peaks is revealing. The closer sites, < 100 nt away from the CLIP peaks, have a relaxed structural requirement compared to more distant sites (Fig. 5B). Because these distance effects should not impact local RNA structure, they suggest that RNA looping contributes quantitatively to the Kd of ADARcd binding (Figure 6; see Discussion) and that local structure serves to localize the ADARcd moiety rather than only for dsRBD binding or for editing site selection (Stephens et al. 2004; Eggington et al. 2011). In other words, editing sites that are closer to the Hrp48 binding sites are more efficiently edited and can tolerate weaker ADARcd binding than sites that are further away. The recent identification of a conserved RNA-binding loop within hADAR2cd (Matthews et al. 2016) also indicates that the ADARcd normally complements the dsRBDs and contributes to substrate binding, perhaps by stabilizing the RNA-protein complex.

**Figure 6.**
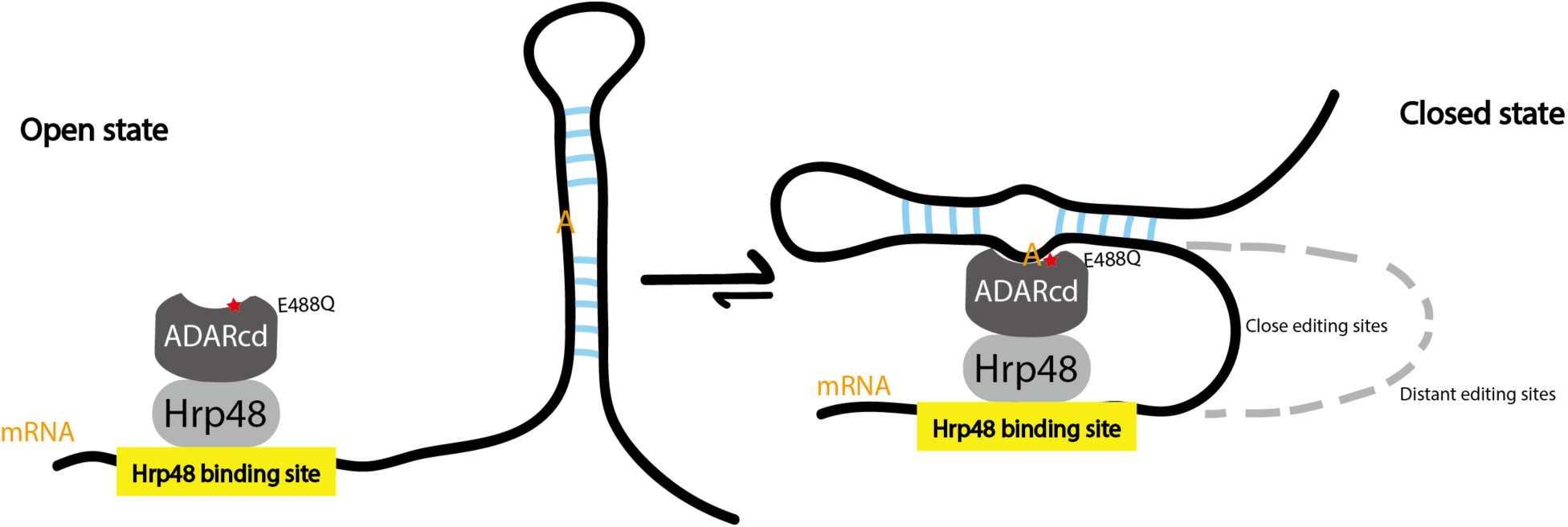
Proposed model: Hyper-TRIBE is better able to shift the equilibrium towards the closed state. When HyperTRIBE binds to a target mRNA, adenosines in potential double-stranded regions are candidate substrates for binding to the HyperADARcd moiety. More proximal adenosines will be accessible for HyperADARcd binding with a small RNA loop, whereas more distant adenosines require a larger and more destabilizing RNA loop. We suggest that the Hyper-ADARcd with its E488Q mutation, is better able to shift the equilibrium toward the closed state, perhaps by binding more tightly to the flipped adenosine. However, the exact mechanism by which the E488Q mutation increases the editing/deamination rate is unknown (Kuttan and Bass 2012).

## Discussion

Our recently developed TRIBE method (McMahon et al. 2016) expresses in vivo a chimeric protein, which is a fusion of a RBP of interest to the catalytic domain of ADAR. TRIBE performs Adenosine-to-Inosine editing on RNAs recognized by the RNA binding moiety of the protein, i.e., the RBP. However, TRIBE-mediated editing is quite selective and probably gives rise to a high false negative rate for identification of RBP target RNAs. We present here the *in vivo* characterization of HyperTRIBE, which carries the hyperactive E488Q mutation within the ADARcd.

The HyperTRIBE results overlap much more successfully than TRIBE results with our previously published Hrp48 CLIP results. These CLIP data were very similar whether from endogenous Hrp48 or from overexpressed Hrp48 TRIBE fusion protein (McMahon et al. 2016), indicating that the TRIBE protein interacts with RNA similarly to endogenous Hrp48. The much better overlap suggests that HyperTRIBE has a much reduced false-negative problem compared to TRIBE. A different HyperTRIBE fusion protein, containing the RBP FMRP, also has more editing sites and genes compared to regular TRIBE with this RBP (McMahon et al. 2016) (Fig. S4). Importantly, perfect overlap with CLIP data is not necessarily expected, i.e., iCLIP data may identify its own set of false positives, meaning RNAs of little interest. They could for example reflect low affinity, transient interactions with transcripts that cross-link efficiently to the RBP.

Might there be a comparable source of HyperTRIBE false positives? Activity by the HyperADARcd alone is a likely candidate, and we are aware of editing by the human HyperADAR1-cd in other systems (Wang et al. 2015). However, we never obtained any substantial editing with expression of only the *Drosophila* HyperADARcd (e.g., Fig. 1A), despite comparable expression levels to the HyperTRIBE fusion protein (data not shown).

A conservative threshold of 10% editing was initially established to ensure that the TRIBE assay was not impacted by substantial numbers of false positives (McMahon et al. 2016). We began this HyperTRIBE study with the same 10% threshold, so that the new data could be directly compared with our previous TRIBE data. Strikingly, HyperTRIBE identifies many more sites and genes above this threshold, consistent with the original characterization of the E488Q mutation in yeast (Kuttan and Bass 2012). More surprising was the realization that many of the new HyperTRIBE > 10% editing sites correspond to proper TRIBE editing sites but with editing frequencies well below the 10% threshold. The correspondence of these sites not only with HyperTRIBE sites but between replicate TRIBE experiments further indicates that they are genuine editing sites. Three conclusions from these data are: 1) HyperTRIBE edits more efficiently than TRIBE; 2) Even very low editing frequencies identify true TRIBE editing sites if they are identified in replicate experiments; 3) there are many more bona fide TRIBE editing events than are identified with conservative thresholds. Nonetheless, many TRIBE sites will require substantial sequencing depth to be identified. Much of this increased sequencing can probably be avoided by using HyperTRIBE, which should more generally be a superior approach for the identification of RBP targets in mammalian as well as fly cells and neurons.

Nonetheless, the fraction of edited adenosines is low even for HyperTRIBE, presumably still reflecting the sequence and structural requirements of the ADARcd, i.e., the nearest neighbor sequence preference and the double-stranded character surrounding a bulged adenosine. This structural landscape is very different from that of random 3’UTR sequences (Fig. 4B), suggesting that the ability to form a local intramolecular helix is important for HyperTRIBE as well as for TRIBE. Nonetheless, HyperTRIBE has a clear reduction in these sequence and structural requirements (Fig. 4A, 4B). The relative lack of structure on the 3’ side of the HyperTRIBE and the below-threshold editing sites is particularly striking (Fig. 4B). The asymmetry is probably less important for HyperTRIBE editing, which may reflect differences in rate-limiting steps, e.g., the ADARcd binding affinity/off-rate is more important for the slower TRIBE enzyme (see below).

The distance effects (Fig. 5A) are presumably a simple consequence of RNA looping between the RBP binding site and the edited region. Similar to the effect of increased loop size on the stability of a RNA hairpin, the stability of a weak protein-RNA interaction should be negatively impacted by a bigger loop size, i.e., a greater distance between the RBP binding site and the RNA editing substrate region. Taken together with the relaxed structural constraints on editing sites close to the Hrp48 binding sites (Fig. 5B), the distance effects suggest that the binding affinity of the ADARcd to the substrate region impacts editing efficiency.

Importantly, looping provides a large positive effect, by increasing the local substrate concentration and overcoming an RNA substrate-ADARcd interaction that is otherwise too weak to generate substantial numbers of editing events in vivo (Fig. 6). The RBP moiety should provide all of the mRNA specificity and almost all of the RNA affinity, for HyperTRIBE as well as for TRIBE, whereas the ADARcd and its binding specificity determine the precise mRNA regions and adenosine(s) that will be edited. The HyperTRIBE data more generally indicate that the broad enhancement of editing frequency by the E488Q mutation can be successfully divorced from substrate binding; this enhancement probably requires that the ADARcd-adenosine interaction is thermodynamically supported by a much more stable RBP-mRNA interaction. A long dwell time of the RBP can also allow the ADARcd to sample different mRNA regions and even different RNA configurations, i.e., the local helix surrounding the substrate adenosine (Fig. 6) may be in dynamic equilibrium with less structured configurations but briefly stabilized by binding of the ADARcd.

As previously discussed (Kuttan and Bass 2012; Matthews et al. 2016), the E488Q mutation either enhances base flipping, perhaps by enhanced amino acid insertion by the HyperTRIBE ADARcd into the RNA A helix, or it has a much higher rate of successful editing/flipping event. Notably, the ADARcd loop that occupies the displaced A during flipping (Matthews et al. 2016) contains amino acid 488. Better binding of the HyperTRIBE ADARcd to its substrate region might enhance the transition state lifetime and therefore editing efficiency (Kuttan and Bass 2012; Matthews et al. 2016). This is further suggested by the impact of distance from the CLIP sites on editing efficiency as well as on the structural requirement surrounding the edited adenosine (Fig. 5). A positive relationship between editing efficiency and substrate binding is also indicated by the enhanced structural requirement of TRIBE sites as compared to below-threshold or HyperTRIBE sites (Fig. 4B). Indeed, we speculate that enhanced editing of the common sites by TRIBE, which is comparable to the editing of the HyperTRIBE unique sites (Fig. 3A), is achieved by better TRIBE binding to these common sites because of the increased structure on the 3’ side of these edited adenosines (Fig. 4B).

## Methods

### Molecular Biology

RBP-ADARcd with E488Q mutation was created by performing Quikchange® Site-directed Mutagenesis on pMT-RBP-ADARcd-V5 plasmid (McMahon et al. 2016). Primers 5’-TCGAGTCCGGTCAGGGGACGATTCC and 5’-GGAATCGTCCCCTGACCGGACTCGA were used to induce point mutation to the underlined nucleotide. 15 AA, 50 AA and 100 AA flexible linkers and 15 AA, 50 AA rigid linkers (Amet et al. 2009) were chemically synthesized by Integrated DNA Technologies, Inc. and cloned into pMT-Hrp48-ADARcd-E488Q-V5 plasmid using Gibson Assembly® from NEB. The other linkers were created by PCR duplicating the fragment and cloning. Transient expression of TRIBE constructs was performed by co-transfecting pMT TRIBE plasmids with pActin-EGFP to *Drosophila* S2 cells using Cellfectin® II from Thermo Fisher Scientific. Cells were allowed 48 hours after transfection for adequate expression of GFP before sorting with BD FACSAria™ II machine for GFP positive cells. Total RNA was extracted from the sorted cells with TRIzol™ LS reagent. TRIBE protein expression was induced with copper sulfate 24 hours before FACS sorting. Expression of all fusion proteins was assayed by transient expression in S2 cells and western blot against V5 tag (Invitrogen, 46-1157). TRIBE stable cell lines used in this paper are the same as the original TRIBE paper (McMahon et al. 2016). GFP labeled neurons are manually selected by using a glass micro-pipette from dissected, digested and triturated fly brains as previously described (Abruzzi et al. 2015; McMahon et al. 2016).

Standard Illumina TruSeq® RNA library Kit was used to construct RNA-seq library from S2 cells. Manually-sorted cells are subjected to RNA-seq library protocol as previously decribed (Abruzzi et al. 2015; McMahon et al. 2016). All libraries were sequencing by Illumina NextSeq® 500 sequencing system using NextSeq® High Output Kit v2 (75 cycles). Each sample were covered by ~20 million raw reads.

### RNA-editing Analysis

The criteria for RNA editing events were: 1) The nucleotide is covered by a minimum of 20 reads in each replicate; 2) More than 80% of genomic DNA reads at this nucleotide is A with zero G (use the reverse complement if annotated gene is in the reverse strand); 3) A minimum of 10% G is observed at this site in mRNA (or C for the reverse strand). Genomic DNA of S2 cell and background fly strain are sequenced to identify and exclude possible polymorphism on DNA level. RNA sequencing data were analyzed as previously described (Rodriguez et al. 2012; McMahon et al. 2016), with minor modifications. Background editing sites found in samples expressing Hyper-ADARcd alone were subtracted from the TRIBE identified editing sites both in S2 cells and in fly neurons. Overlap of editing sites from two datasets was identified using “bedtools intersect” with parameters “-f 0.9 -r”.

Quantification of RNA sequencing reads distribution was performed with read_distribution.py script in RSeQC v2.3.7 (Wang et al. 2012).

RNA structure folding analysis was carried out with UNAFold (Markham and Zuker 2008) on flanking sequences of Hrp48 TRIBE editing sites, Hyper TRIBE editing sites, TRIBE below-threshold editing sites and random 3’ UTR sites centered around an “A”, which excludes any genes with Hrp48 CLIP or Hyper TRIBE signal. One thousand sites were blindly selected in each pool, except 221 sites were selected from TRIBE edit sites due its small size. In order to remove oversampling of any region, only one site is considered per gene. A flanking region of 250nt both 5’ and 3’ of the editing site or random site was folded with UNAFold parameters (hybrid-ss-min --suffix DAT --mfold=5,8,200 --noisolate). Base pairing was counted in the predicted minimum free energy (MFE) and predicted suboptimal structures. A profile of double-strandedness is created for each sequence, which is then averaged over all 500 sequences and plotted.

### Fly lines

HyperTRIBE injection plasmid were generated by site-directed mutagenesis of pJFRC7-20xUAS-Hrp48-ADARcd-V5 plasmid to introduce the E488Q mutation. The Hyper-ADARcd only construct is generated in the same way. The transgenes were injected by Rainbow Transgenic Flies, Inc. (Camarillo, CA). UAS-RBP-ADARcd-V5; UAS-eGFP flies (Bloomington stock center #1522) were crossed to elav-gsg-Gal4 driver line to allow adult-only expression of the fusion proteins in all neurons due to the lethality of constitutive pan-neuronal expression (Osterwalder et al. 2001; McMahon et al. 2016). Prior to manual cell sorting, food containing RU486 (0.2 μg/ml, Sigma) was used to induce transgene expression in young flies (~3 days old) for 3 days.

### Data availability

The accession number for the raw sequencing data and processed RNA-editing tracks reported in this paper is NCBI GEO: GSE102814. The scripts used in the analysis are available on GitHub (<u>https://github.com/rosbashlab/TRIBE</u>).

## Acknowledgements

Special thanks are due to Aoife McMahon, a prior lab member, for her important input into this work. We thank Joshua Rosenthal, Brenda Bass, Amy Lee and Aoife McMahon for helpful comments on a very early version of this manuscript as well as current Rosbash lab members for comments and discussion. The work was supported by the Howard Hughes Medical Institute and by a NIH EUREKA grant (DA037721).

## Author Contributions

Conceptualization, W.X., M.R. and Aoife McMahon; Investigation, W.X.; Software, R.R.; Formal Analysis, W.X. and R.R.; Visualization, W.X. and R.R.; Writing – Original Draft, W.X. and M.R.; Funding Acquisition, M.R.

---

**Figure S1.**
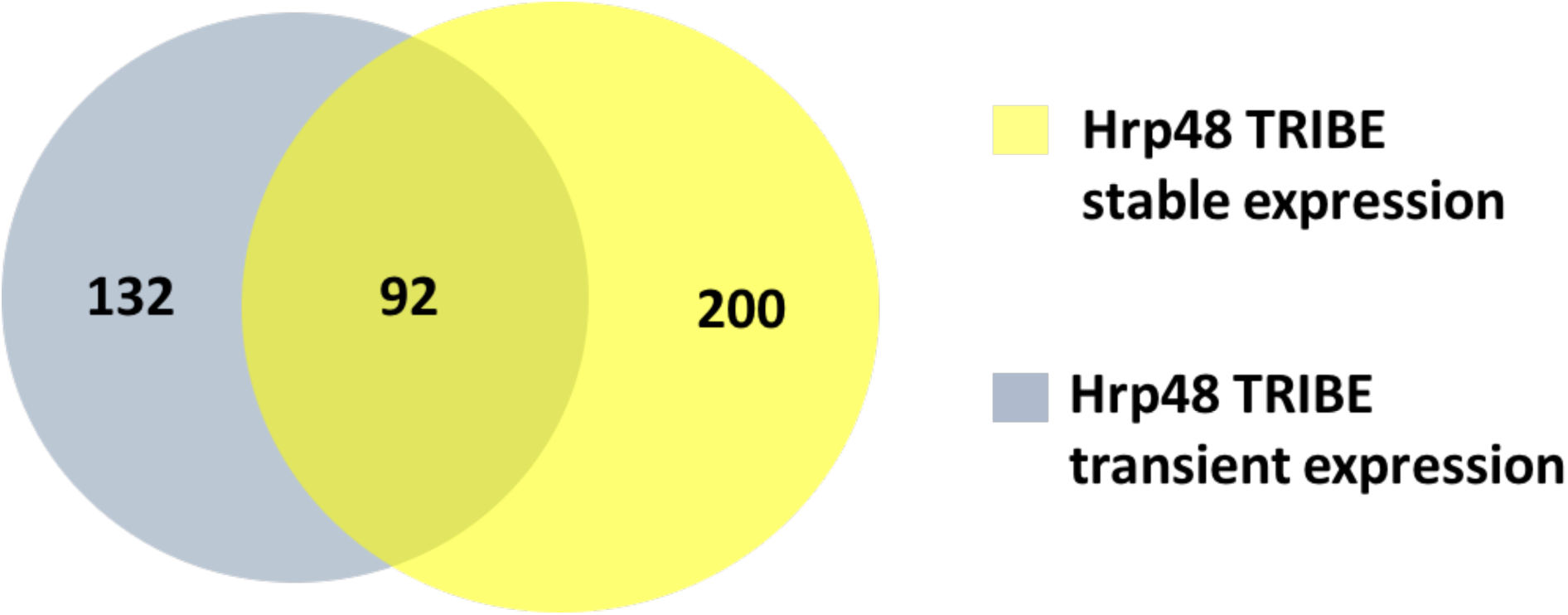
Comparison of Hrp48 TRIBE target genes between stable and transient expression. Venn diagram shows the overlap of genes identified by expressing TRIBE in S2 cells transiently (224 in total, blue) and stably (292 in total yellow).

**Figure S2.**
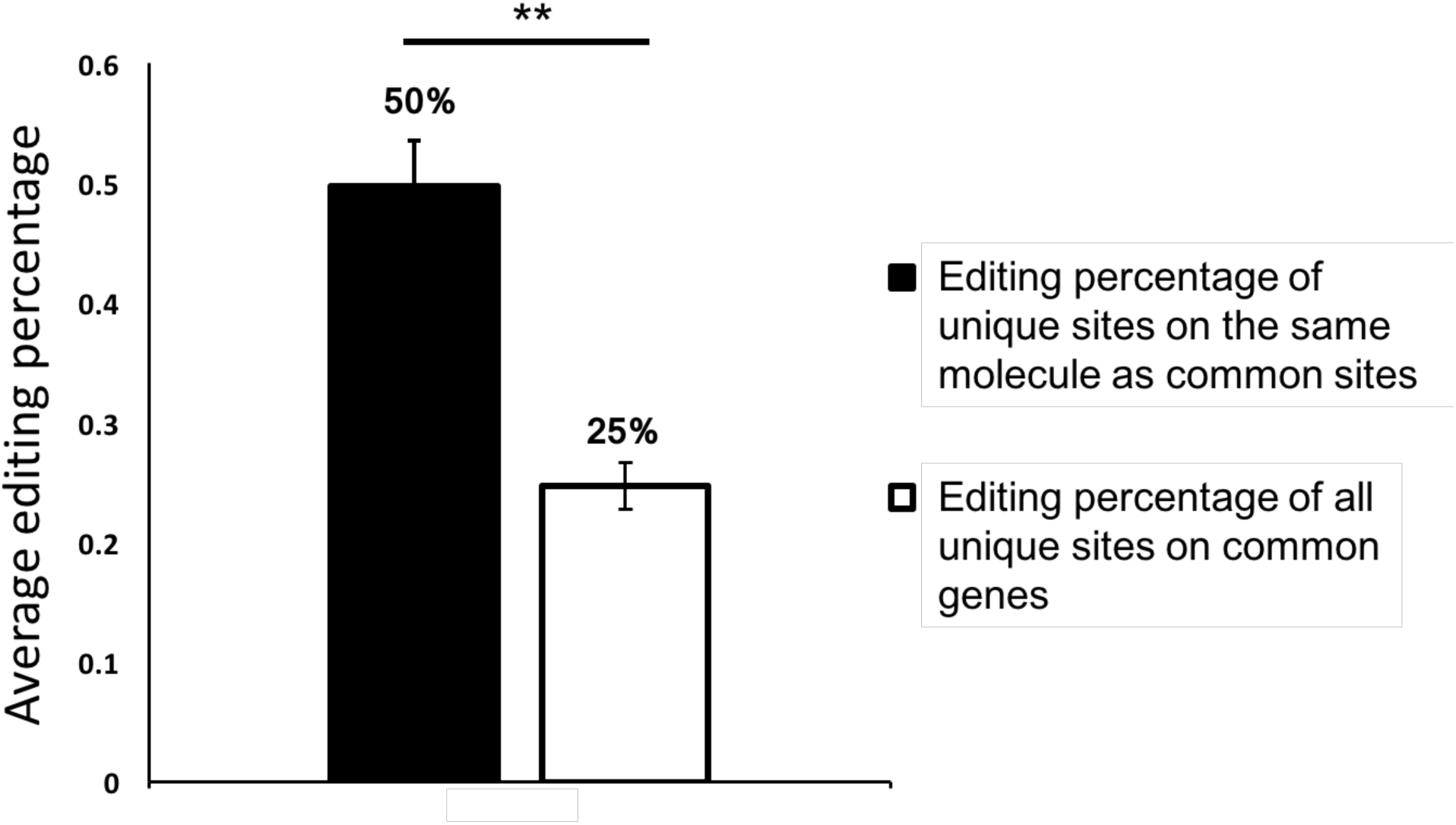
Multiple editing events on the same molecule are positively correlated. Bar graph showing the percentage of the adenosines edited under different circumstances. Results are summarized by manually browsing through mapped sequence reads of the HyperTRIBE sample in IGV. The editing percentage of total unique editing sites on genes that are commonly identified by TRIBE and HyperTRIBE are calculated (white bar, 25%), and the editing percentage of the unique sites when the common sites are edited on the same molecule are calculated (black bar, 50%). When the common sites are edited on the same molecule, the other sites are edited at much higher frequency (Student t-test performed, p<1E-09), indicating multiple editing events on the same molecule are not independent.

**Figure S3.**
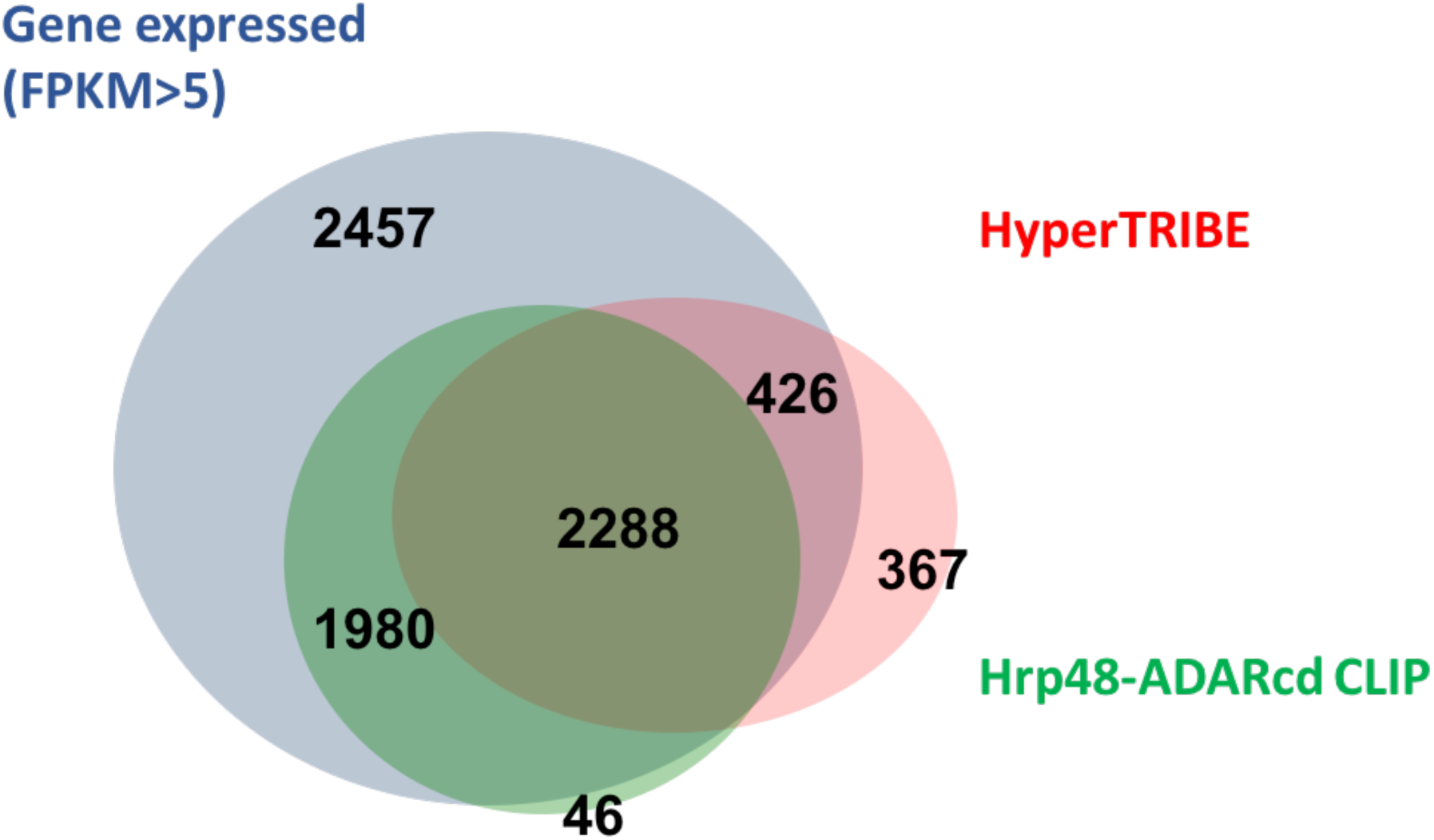
Genes identified by HyperTRIBE overlap well with CLIP-identified genes. Venn diagram shows the overlap of genes identified by HyperTRIBE (3085 in total, pink), Hrp48-ADARcd CLIP (4327 in total, green) and all expressed genes in the cells (7151 in total, blue). HyperTRIBE-identified genes are significantly enriched for CLIP-positive genes (Z-test performed, p<0.0001, ratio of overlap with HyperTRIBE genes for CLIP-positive and CLIP-negative genes compared), indicating specificity.

**Figure S4.**
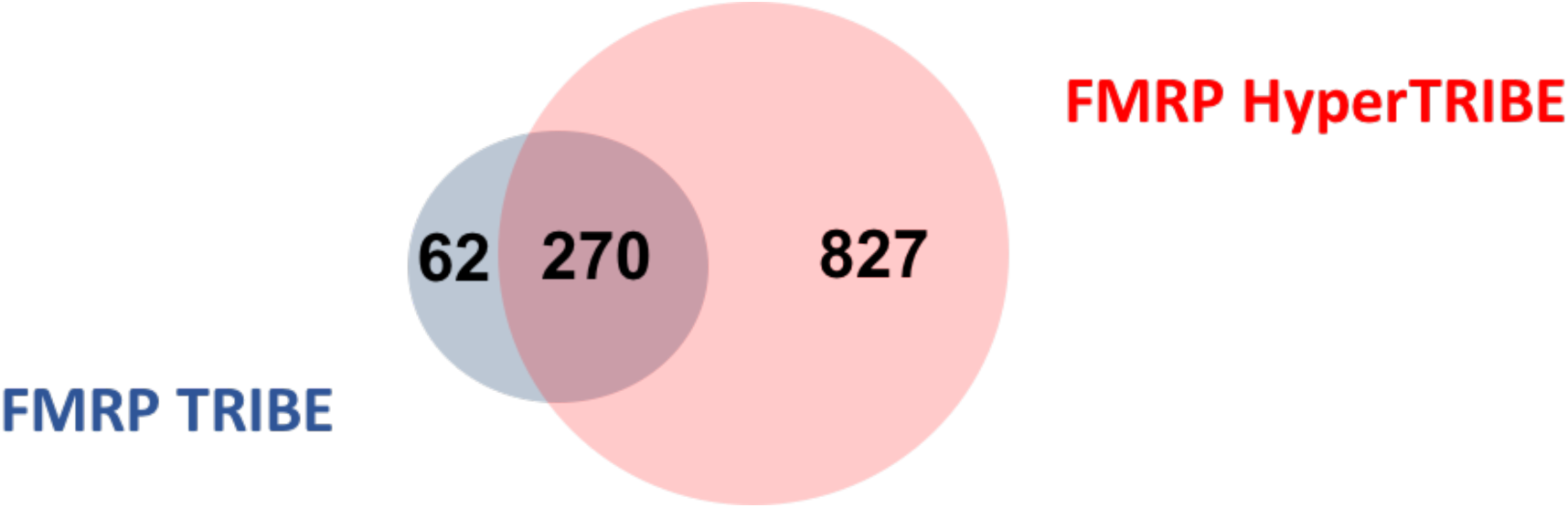
When fused to FMRP, HyperTRIBE also increases editing efficiency with high specificity compared to TRIBE. Venn diagram showing the overlap of genes identified by FMRP HyperTRIBE and TRIBE when both are transiently expressed in S2 cells. FMRP HyperTRIBE-targeted genes (1097 in total, pink) overlap well with FMRP TRIBE (332 in total, blue)-target genes, but there are three times as many HyperTRIBE genes.

**Figure S5.**
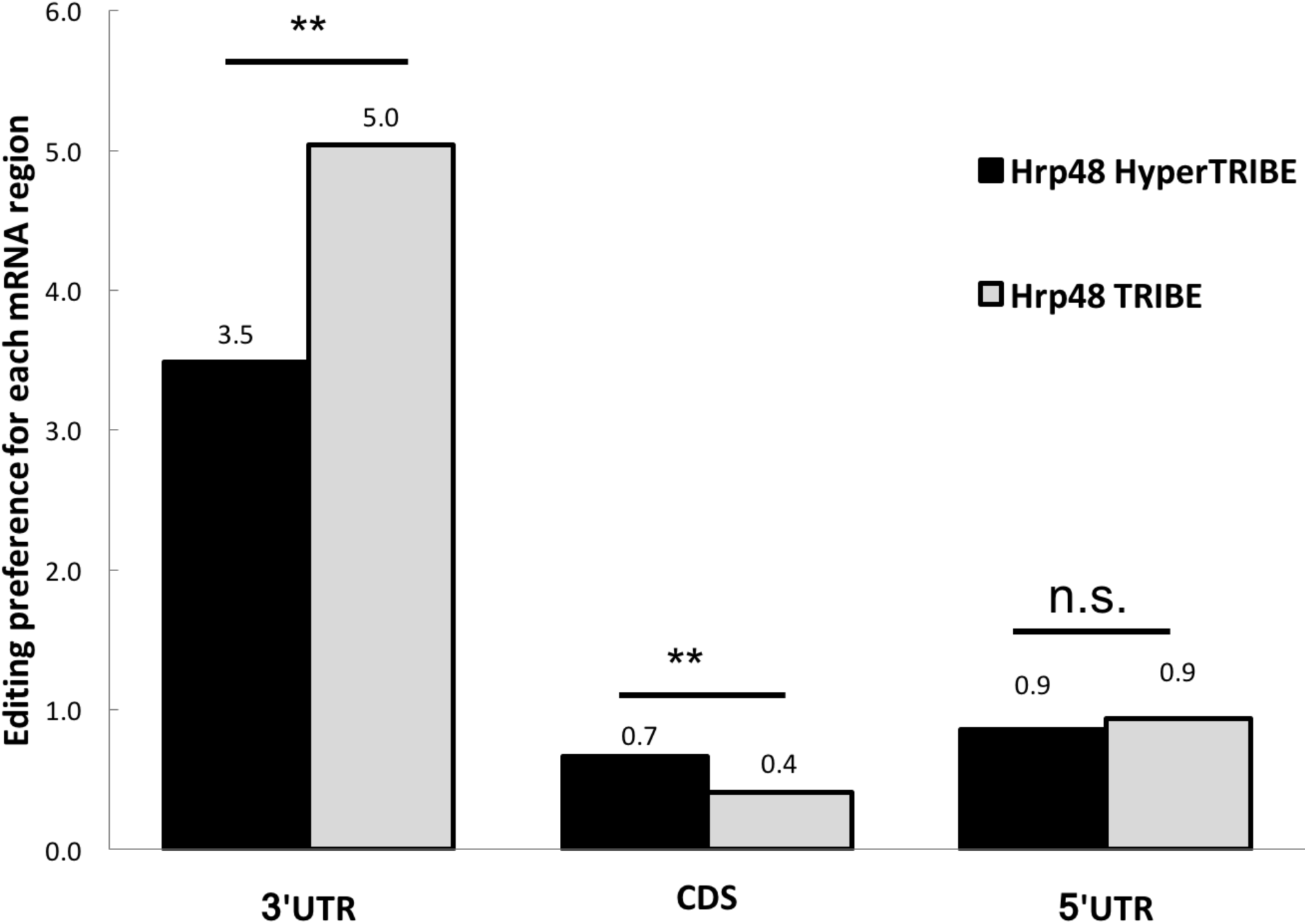
HyperTRIBE editing retains the 3’ UTR binding preference of Hrp48 protein. Both TRIBE methods show 3’UTR editing preference, although the preference is lower for HyperTRIBE. Editing preference is calculated as the distribution of editing sites in each mRNA region normalized to the distribution of sequence reads in each mRNA region. (n.s.=p>0.05, **p<0.001, Z-test performed HyperTRIBE compared against TRIBE)

